# Impact of single mutations on binding kinetics of triplex forming oligos revealed by fluorescence proximity sensing in heliX^®^ biosensor

**DOI:** 10.1101/2022.11.21.517309

**Authors:** Vera Molkenthin, David Baumstark, Thomas Schubert, Gernot Längst, Maximilian G. Plach

**Affiliations:** 2bind GmbH, Regensburg, Germany; Faculty of Biology and Preclinical Medicine, University of Regensburg, Germany

**Author notes:** 2bind.com.

## Abstract

The sequence specific association of RNA with DNA via formation of RNA/DNA triple helices is relevant for regulation of gene expression, repair mechanisms, and chromatin organization. Formation of such RNA/DNA triplexes requires poly-purine sequences, which allow for Hoogsteen base-paring in addition to Watson-Crick pairing in the major groove of DNA. Binding of triplex forming oligos (TFOs) is sequence-specific and understanding sequence dependencies is key for the development of targeted tools for gene therapy. Here, we highlight a direct approach for determining binding kinetics and binding constants for TFOs using the state-of-the-art heliX®biosensor tool. With this, we provide key insights into the binding kinetics of RNA and DNA TFOs to a triplex targeting site (TTS)-containing DNA double helix measured in real-time. Dependent on the introduced base, point mutations in one position of a triplex forming oligo (TFO) can change the dissociation constant (KD) by several orders of magnitude or just by one log, affecting primarily the dissociation rate. Furthermore, we demonstrate that the heliX®biosensor assay is also well-suited for detection of rather weak triplex formation. The weakest binding we could identify was 140 μM, for a TFO, which other studies considered as non-binding.

## Introduction

The well known structure of the DNA double helix leaves room for a third strand of nucleic acids in its major groove. A third nucleotide can attach to a Watson-Crick base pair via Hoogsteen base-pairing. The possibility of DNA to therefore form triple helices is known since half a century [1, 2, 3]. RNA/DNA triplexes occur in nature and are one of the mechanisms that regulate gene expression [4, 5, 6, 7, 8, 9]. TFOs have been recognized as tools for gene manipulation [10, 11, 12, 13] and RNA/DNA triple helices have raised great interest as therapeutic targets [14].

### Triplex structure and formation

RNA or DNA TFOs bind to the major groove of duplex DNA. Formation of the RNA/DNA or DNA triplexes requires that one strand of the double helix contains several purines in a row [2, 15, 16]. Such poly-purine stretches can be bound by three different types of TFOs: (i) pyrimidine TFOs, (ii) purine TFOs and [iii] purine-pyrimidine TFOs [16]. Formation of triple helices requires the presence of divalent cations and is sensitive to other buffer conditions, like pH, salt concentration, and detergent content [1, 16, 17]. Previously, interactions between TFOs and DNA double helices containing TTSs have been studied by electrophoretic mobility shift assays, Isothermal Titration Calorimetry (ITC), melting temperature analysis, Microscale Thermophoresis (MST), molecular beacon strategies, nuclease digestions, ultraviolet absorbance decay measurements [17, 16, 18, 19, 20, 21, 22], as well as real time biosensor studies [23]. Frequently, triple helices have been considered as unstable and transient structures, which hardly form under physiological conditions [13]. However, MST studies revealed that binding affinities of TFOs to TTSs are in the ranges of specific DNA-protein interactions [17]. Biosensors enable the measurement of real time kinetics and offer further insight into the dynamics of molecular interactions. The heliX®biosensor offers the unique possibility to immobilize nucleotide sequences via hybridization on the surface and is the ideal tool to study interactions of nucleic acids in real time.

### heliX®biosensor

heliX®biosensors enable the immobilization of target nucleotide strands via hybridization Figure 2. The heliX®biosensor adapter chips are coated with single stranded DNA anchors on the surface. Universally usable adapter strands carry a fluorophore and hybridize to the anchors on the chip, leaving a 48 nt single stranded overhang. For immobilization by hybridization, the sequence of interest is prolonged by a sequence that is the reverse complement of the 48 nt single stranded overhang. The technology is frequently used for DNA-targeted immobilization of DNA-protein conjugates [24]. The orientation of the negatively charged nanolever can be influenced by applied voltage, a technology called switchSENSE^®^, which has been developed by Dynamic Biosensors (DBS), Munich, Germany [25]. The heliX®biosensor offers two ways to detect binding of analytes to the molecule of interest: (i) Monitoring the influence of the binding event on the switching speed of DNA nanolevers in the dynamic measurement mode with alternating voltage, and (ii) Fluorescence Proximity Sensing (FPS), detecting the effect of changes in the fluorophore’s local environment on the fluorescence intensity [26]. FPS is measured in the static measurement mode, with a constant negative voltage, which ensures that the DNA strands are expelled from the electrode and presenting the target of interest in a certain distance from the surface.

**Figure 1.**
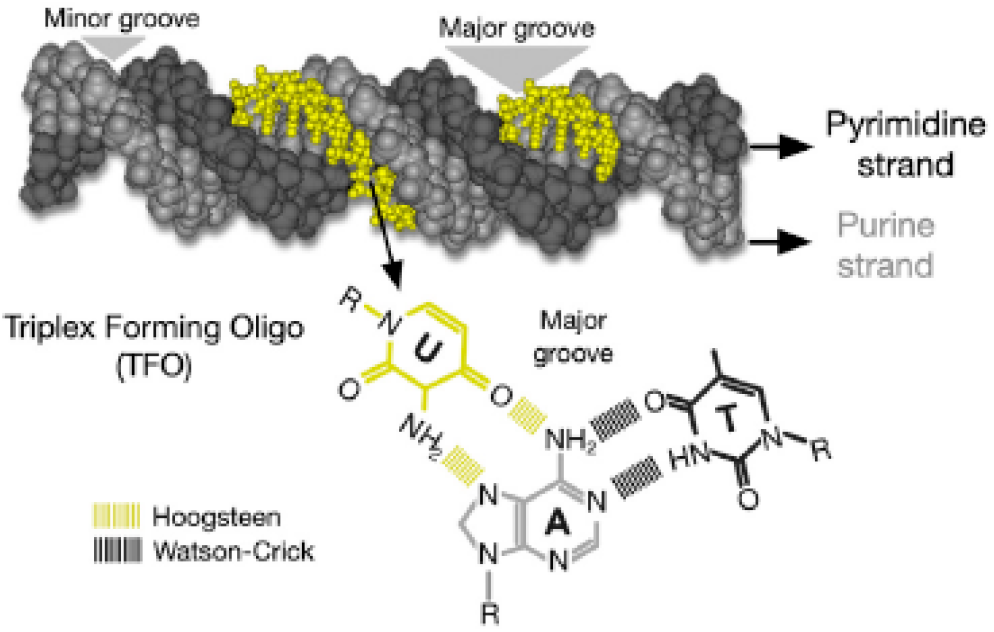
Schematic representation of triplex formation. From [16].

**Figure 2.**
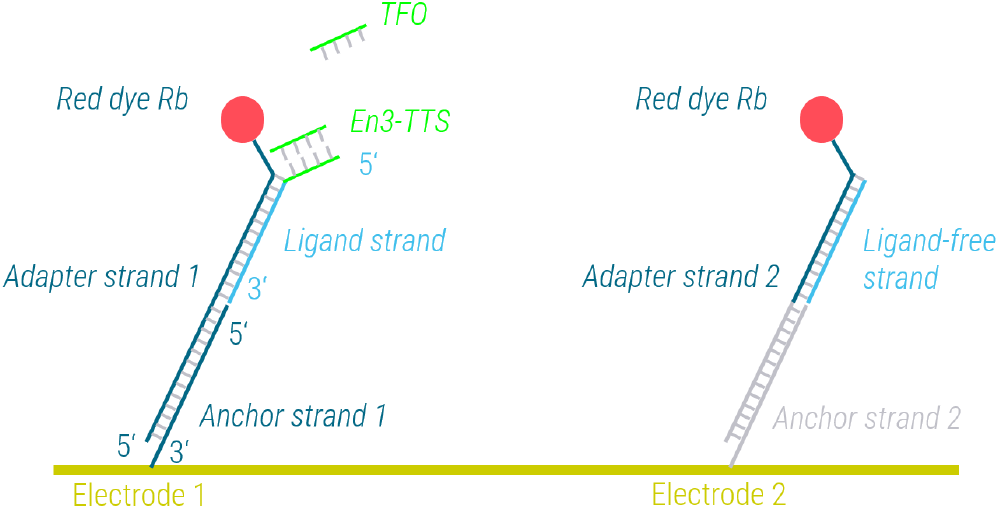
Schematic representation of the assay setup.

### The En3-TTS motif

Maldano et al. [17] discovered the En3-TTS motif using the software Triplexator [27], which predicts sequences with high potential to form triple helices. The En3-TTS motif consists of the symmetric core motif TC(T)_15_C(T)_6_CCTCC. The sequence was prolonged by the reverse complement of the adapter strand and immobilized on the chip. The reverse complement of the En3-TTS motif was added to build the double stranded target sequence. We compared binding of the pyrimidine RNA-TFO CCUCC(U)_6_CU_15_CU and the respective purine RNA-TFO AG(A)_15_G(A)_6_GGAGG to the double helix on the chip. A DNA-TFO and a RNA-TFO with identical sequence were compared. Additionally, we analyzed the impact of changing the C in position 12 to all three alternative bases on the association and dissociation of the TFO.

## Results and Discussion

### Fluorphores exhibit different sensitivity for DNA in FPS

The six different dyes offered by DBS differ largely in their ability to sense the hybridization of the reverse complement and the single stranded DNA sequence overhang immobilized on the chip surface Figure 5A. The largest amplitude is measured with dye Rb, followed by dye Gc. The superiority of dye Rb compared to dye Ra is obvious in binding kinetics of En3-pyrimidine (Y)-DNA-TFO binding to the double stranded En3-TTS motif immobilized on the chip as well Figure 5B. All further experiments were carried out with dye Rb.

### Influence of base family and sugar residue on the binding of TFOs

Binding of the En3-Y-RNA-TFO is very well detectable in the described setup using the heliX®biosensor (Figure 5C). Even the lowest tested concentration of 41 nM results in a detectable signal. An affinity of 7 nM was calculated from three repeated measurements for En3-Y-RNA-TFO (Table 1). In contrast, binding of the En3-purine (R)-RNA-TFO is hardly detectable and the obtained signals are not usable for a kinetic evaluation. A En3-Y-DNA-TFO with identical sequence compared to the En3-Y-RNA-TFO binds with 9 nM affinity (Table 1). Although KDs are in the same range for both TFOs, the binding reaction is in general faster for En3-Y-DNA-TFO, with a two-fold faster association and a two-fold faster dissociation rate (Figure 4). Additionally, the En3-Y-DNA-TFO shows a biphasic dissociation, with an initial phase of fast dissociation. Potentially, En3-Y-RNA-TFOs exhibit such a fast dissociation phase as well, but the contribution of the first dissociation phase to the overall dissociation amplitude is neglectable (Figure 3). Surprisingly, chip saturation was reached at lower amplitudes for En3-Y-DNA-TFO compared to the En3-Y-RNA-TFO (Figure 5C). Further investigations should clarify, whether this is due to a different effect of DNA and RNA on the dye, or whether indeed a lower number of binding sites is accessible for binding of DNA-TFOs compared to RNA-TFOs.

**Table 1.** Calculated kinetic parameters.

| TFO | ka (M<sup>-1</sup>s<sup>-1</sup>) | kd1 (s<sup>-1</sup>) | kd2 (s<sup>-1</sup>) | dissociation amplitude 1 | dissociation amplitude 2 | KD (M) | KD (M) MST<sup>a</sup> |
| --- | --- | --- | --- | --- | --- | --- | --- |
| En3-R-RNA-TFO |  |  |  |  |  | NA | no binding detected |
| En3-Y-RNA-TFO | 1.28E+04 | 6.58E-03 | 9.07E-05 | 0.10 | 0.93 | 7.10E-09 | 1.13E-08 |
| En3-Y-DNA-TFO C12T | 9.09E+02 | 1.27E-01 | NA | 1.00 | NA | 1.40E-04 | no binding detected |
| En3-Y-DNA-TFO C12A | 1.69E+04 | 1.84E-02 | 2.75E-03 | 0.53 | 0.47 | 1.65E-07 | 2.83E-07 |
| En3-Y-DNA-TFO C12G | 2.01E+04 | 8.66E-03 | 1.54E-03 | 0.68 | 0.32 | 7.60E-08 | comparable to C12A |
| En3-Y-DNA-TFO | 2.43E+04 | 1.72E-01 | 1.99E-04 | 0.08 | 0.92 | 8.76E-09 | 1.63E-08 |

<sup>a</sup> from [17]

**Figure 3.**
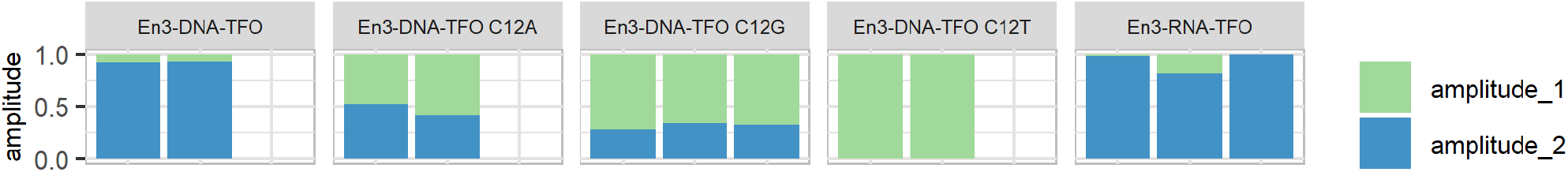
Dissociation amplitudes of phase 1 and phase 2 of biphasic TFO dissociation. Most measured TFOs show biphasic binding behaviour. The two phases of dissociation show different extent of contribution to the overall dissociation.

**Figure 4.**
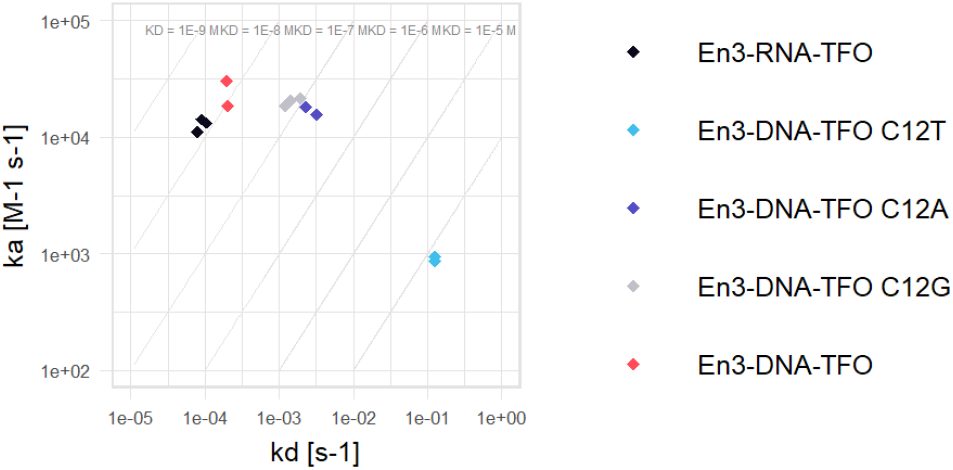
kon-koff rate map of measured interactions. Association and dissociation rates are plotted on y- and x-axis, respectively. The values were calculated from two to three experiments per TFO, each measured in five serial dilutions. In the case of biphasic dissociations, kd2 values were included in the figure.

### Influence of point mutations on the binding of pyrimidine DNA-TFOs

Single point mutations in position C12 change the kinetic binding behaviour of En3-Y-DNA-TFOs dramatically (Figure 5D). All four interactions show biphasic dissociation rates. In case of the two En3-Y-DNA-TFOs with purines in position 12 (C12A and C12G), the two dissociation rates differ by less than factor 10 (Figure 5D), and contribute more or less equally to the total amplitude of dissociation (Figure 3). Dissociation of the two En3-Y-DNA-TFOs with pyrimidines in position 12 (unchanged and C12T) is defined by two dissociation phases with different dissociation rates and amplitudes. The unchanged En3-Y-DNA-TFO shows a fast first dissociation phase with minor contribution to the total amplitude, followed by a slow second dissociation phase. In the case of C12T the fast first dissociation phase is the major contributor to the amplitude of dissociation. The second dissociation rate of C12T cannot be determined within the recorded time frame, but part of the material appears to remain on the surface or dissociates very slowly. Potentially, comparisons of remaining material of unchanged En3-Y-DNA-TFO and C12T after longer dissociation times should be considered as additional criterion to fully characterize the binding behaviour of the two TFOs. In general, dissociation rates are more affected by mutations than association rates (Figure 4). C12A and C12G mutations increased the KD value by one order of magnitude, whereas the C12T mutation resulted in an increase by four orders of magnitude.

**Figure 5.**
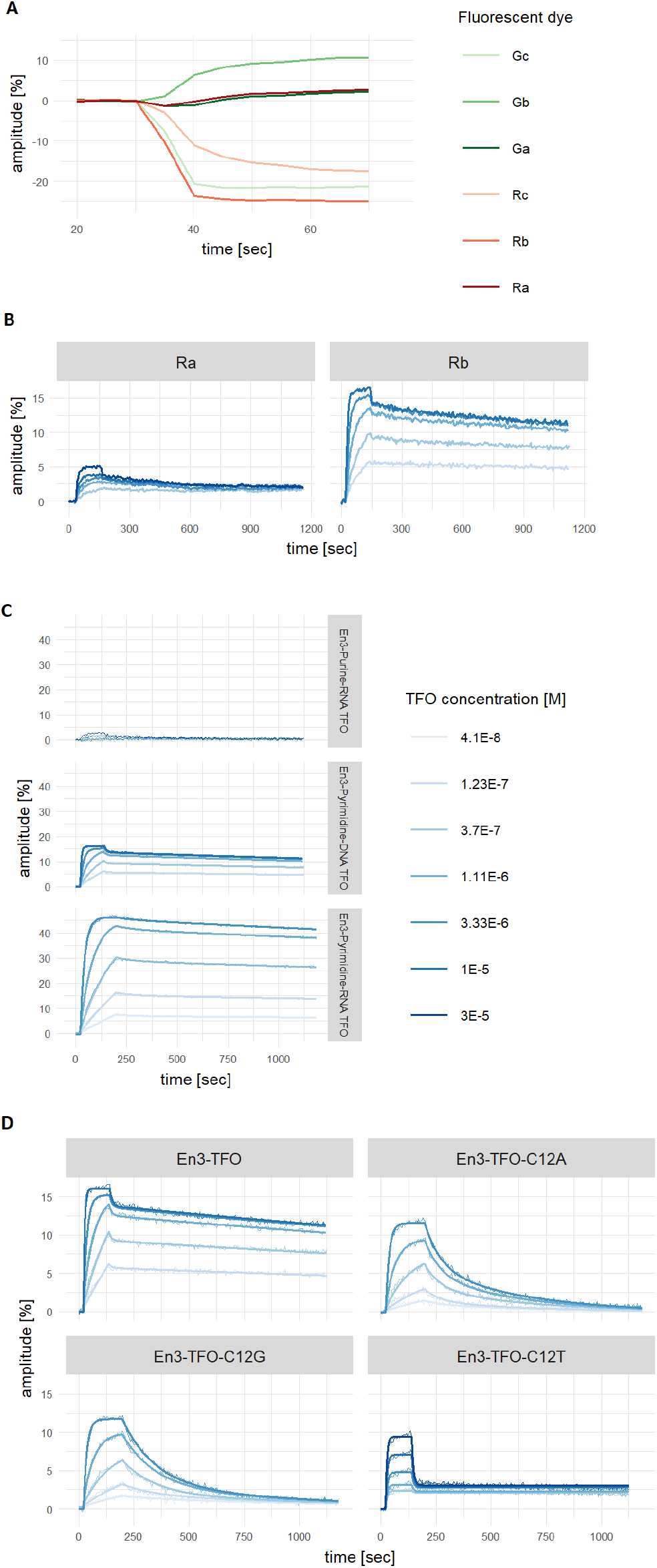
Dye scouting, influence of base family, sugar residue and point mutations on binding of TFOs. (A) Binding of the reverse complement in 1E-7 M concentration to the En3-TTS motif immobilized on the chip was probed with six different dyes from DBS. (B) Binding kinetics of En3-Y-DNA-TFO were measured with dye Ra and dye Rb in comparison. (C) Binding of DNA/RNA, pyrimidine/purine TFOs to immoblized double stranded En3-TTS motif. (D) Binding of different DNA-TFOs to immoblized double stranded En3-TTS motif. TFOs were added in indicated concentration in TA buffer (40 mM Tris-acetate (pH7.4), 10 mM Mg-acetate, 0.05% NP40). Assay temperature: 25°C. Plots include double referenced data and curve fits.

### Comparison of heliX®biosensor and MST results

We used the same model system as studied previously with MST [17], and had the chance to compare affinities of molecular interaction determined with two independent methods. MST measures interactions of molecules in solution. One binding partner is labeled with a fluorophore and the second binding partner is titrated. Binding events affect the temperature related intensity change (TRIC) of the fluorophore and thereby KD values can be determined from relative fluorescence values plotted against the concentration of the unmodified binding partner. In contrast to MST in the heliX®biosensor assay one of the binding partners is anchored to the biosensor chip surface, but still distant from the surface due to the switchSENSE®technology. Furthermore, in the heliX®biosensor assay affinities are calculated from association and dissociation rates, whereas MST is an equilibrium based method for affinity determination. Despite the completely different measurement principles, the KD values determined in both assays correspond very well [17] (Table 1). Values differ by less than factor 2 and, in general, the heliX®biosensor assay determined slightly lower KD values. Interestingly, C12T did not show any detectable interaction in MST, whereas binding was detectable in the heliX®biosensor assay, albeit with a very high KD value of 140 μM. Equilibrium methods require high concentrations of the unlabeled binding partner to determine KD values for weak interactions. For practical reasons, the required concentrations might not have been reached in the previous MST study.

## Conclusions

En3-TFOs designed in Maldonado et al. [17] bind with affinities of 140 *μ*M to 7 nM and exhibit major differences in the dissociation rates. The determined KD values reflect the previous findings very well [17] and provide further evidence that interactions of TFOs and a DNA double helix can reach affinities in the same range as affinities reported for protein-protein or protein-DNA interactions. The setup of heliX®biosensor assays for measuring the interaction of nucleic acids is very straight forward and takes advantage of the possibility to immobilize nucleic acids via hybridization. Details on binding kinetics are frequently required to fully understand the interaction of two molecules. In particular, the method is considered as highly useful for designing nucleic acids for therapeutic applications.

## Materials and Methods

### Oligonucleotides

HPLC purified oligonucleotides were ordered from Sigma-Aldrich. The En3-TTS-ligand strand was designed by adding the reverse complement of adapter strand 1 (AS-1) to the 3’end of the target sequence, separated by an interspacing triple T motif. Oligonucleotides were dissolved in Tris-EDTA (10 mM Tris-HCl pH8, 0.1 mM EDTA).

### Hybridization of AS-1 and ligand strand

For hybridization 200 nM AS-1 and 250 nM En3-TTS-ligand strand were mixed in TE40 buffer (10 mM Tris-HCl pH7.4, 40 mM NaCl, 0.05 mM EDTA, 0.05 mM EGTA, 0.05 % Tween20,) and incubated for 20 min at 25°C, 600rpm.

### Dye scouting

Dye scouting was carried out using the Dye Scouting Kit of DBS (Cat.#DS-6). The kit contains AS-1 and adapter strand 2 (AS-2) conjugated with six different dyes. AS-2 is prehybridized with ligand-free strand (LFS). Hybridized En3-TTS-ligand strand/AS-1 and hybridized LFS/AS-2 were mixed in equimolar concentration and injected into the adapter biochip (DBS, Cat.#ADP-48-2-0) during the functionalization procedure. Functionalization time was set to 200 sec. To measure the binding, the reverse complement of the En3-TTS motif was injected in 1E-7M concentration in TA buffer (40 mM Tris-acetate pH7.4, 10 mM Mg-acetate, 0.05% NP-40) with a flow rate of 20 *μ*L/min. TA buffer was also used as running buffer. LED power was set to 4%, and green or red LED were selected based on the measured dye. Sampling rate was set to 0.2 Hz.

### Kinetics

Functionalization was carried out as described under the section “Dye scouting”. To build the double stranded En3-TTS, 35 *μ*L of the reverse complement of the En3-TTS motif was injected in a concentration of 1.23E-8 M in TA buffer with a flow rate of 20 *μ*L/min. To determine kinetic values, the TFOs were injected in five serial dilutions in TA buffer with a flow rate of 20 *μ*L/min. Association was followed for 120 or 180 sec. Dissociation flow rate was set to 200 *μ*L/min. Signals measured in spot 1 were referenced with signals measured in spot 2 and a measurement of TA buffer without analyte was subtracted. Double referenced curves were fitted using the heliOS software, using “Kinetics Fit with Free Amplitudes” for C12T and “Monophasic Association-Biphasic Dissociation Fit with Continuous Amplitude” for the remaining interactions.

### Plotting

Data was plotted using R [28].

## References

[1] Felsenfeld G, Davies DR & Rich A. Journal of the American Chemical Society 79(8):2023 (1957)

[2] Hoogsteen K. Acta crystallographica 12(10):822 (1959)

[3] Hoogsteen K. Acta Crystallographica 16(9):907 (1963)

[4] Carbone GM, Napoli S, Valentini A et al. Nucleic acids research 32(14):4358 (2004)

[5] Goñi JR, De La Cruz X & Orozco M. Nucleic acids research 32(1):354 (2004)

[6] Goñi JR, Vaquerizas JM, Dopazo J et al. BMC genomics 7(1):1 (2006)

[7] Mondal T, Subhash S, Vaid R et al. Nature communications 6(1):1 (2015)

[8] Postepska-Igielska A, Giwojna A, Gasri-Plotnitsky L et al. Molecular cell 60(4):626 (2015)

[9] Roy C. Nucleic acids research 21(12):2845 (1993)

[10] Barre FX, Ait-Si-Ali S, Giovannangeli C et al. Proceedings of the National Academy of Sciences 97(7):3084 (2000)

[11] Moser HE & Dervan PB. Science 238(4827):645 (1987)

[12] Le Doan T, Perrouault L, Praseuth D et al. Nucleic acids research 15(19):7749 (1987)

[13] Duca M, Vekhoff P, Oussedik K et al. Nucleic acids research 36(16):5123 (2008)

[14] Shafer RH. Progress in nucleic acid research and molecular biology 59:55 (1997)

[15] Rajagopal P & Feigon J. Biochemistry 28(19):7859 (1989)

[16] Maldonado R, Schwartz U, Silberhorn E et al. Molecular cell 73(6):1243 (2019)

[17] Maldonado R, Filarsky M, Grummt I et al. RNA 24(3):371 (2018)

[18] Tateishi-Karimata H, Nakano M & Sugimoto N. Scientific reports 4(1):1 (2014)

[19] Nakanishi M, Guntaka RV & Weber KT. Nucleic acids research 26(22):5218 (1998)

[20] Xodo LE. European journal of biochemistry 228(3):918 (1995)

[21] Antony T & Subramaniam V. Antisense and Nucleic Acid Drug Development 12(3):145 (2002)

[22] Torigoe H, Hari Y, Sekiguchi M et al. Journal of Biological Chemistry 276(4):2354 (2001)

[23] Wood S. Microchemical journal 47(3):330 (1993)

[24] Schulte C, Soldà A, Spänig S et al. Communications biology 5(1):1 (2022)

[25] Cléry A, Sohier TJ, Welte T et al. Methods 118:137 (2017)

[26] Häußermann K, Young G, Kukura P et al. Angewandte Chemie 131(23):7744 (2019)

[27] Buske FA, Bauer DC, Mattick JS et al. Genome research 22(7):1372 (2012)

[28] R Core Team. R: A Language and Environment for Statistical Computing. R Foundation for Statistical Computing, Vienna, Austria (2022)

